# Perturbation robustness analyses reveal important parameters in variant interpretation pipelines

**DOI:** 10.1101/2020.06.29.173815

**Authors:** Yaqiong Wang, Aashish N. Adhikari, Uma Sunderam, Mark N. Kvale, Robert J. Currier, Renata C. Gallagher, Pui-Yan Kwok, Jennifer M. Puck, Rajgopal Srinivasan, Steven E. Brenner

## Abstract

**Motivation:** Genome sequencing is being used routinely in clinical and research applications, but subsequent variant interpretation pipelines can vary widely. A systematic approach for exploring parameter choices and selection plays an important role in designing robust pipelines for specific clinical applications.

**Results:** We present a framework to be applied in scenarios with limited data whereby expert knowledge informs pipeline refinement. Starting from initial reference variant interpretation pipelines with commonly used parameters, we derived pipelines by perturbing the parameters one by one to determine which parameters can yield meaningful changes in a pipeline’s performance. We updated the reference pipeline by fixing the value of parameters which have small impact on the pipeline’s performance. Then we conducted new rounds of perturbation as the process converged, yielding a stable pipeline which is robust. We applied the framework for genetic disease prediction in de-identified exomes from a cohort of 138 individuals with rare Mendelian inborn errors of metabolism (IEMs) and systematically explored how perturbing different parameters affected the pipeline’s sensitivity and specificity. For this application, we perturbed commonly used parameters in variant interpretation pipelines, including choices of genes, variant callers, transcript models, databases of allele frequencies, databases of curated disease variants, and tools for variant impact prediction. Our analyses showed that choice of variant callers, variant impact prediction tools, MAF threshold, and MAF databases can meaningfully alter results from a pipeline. This work informs the development of exome analysis pipelines designed for newborn metabolic disorder screening and suggests the general application of perturbation analysis in genome interpretation pipeline design.

## 1. Introduction

High throughput sequencing technologies have been adopted in clinical settings in recent years to discover the molecular basis of human diseases. Even though clinical exome/genome sequencing is fairly standard now, subsequent analysis and interpretation of the sequencing data can vary substantially depending on the clinical context and application (Wright, et al., 2018). It is generally understood that parameters in analysis pipelines need to be optimized, but most sequencing studies typically describe only the final set of parameters chosen, without reporting which parameters are most relevant and how altering those parameters would have affected the results.

The analysis of exomes and genomes in rare genetic diseases typically involves sifting through thousands of variants to locate a few that are potentially relevant to patient phenotypes. To design variant interpretation pipelines for this purpose, commonly used parameters may include choice of different variant callers, transcript models, databases of variant frequencies in population, databases of predicted and curated variant impacts, and inheritance models. For some of these parameters, however, numerous choices exist, and there are even more permutations in which these parameters can be combined into an analysis pipeline.

Accurate identification of genetic variants is the foundation for downstream analyses in genomic sequencing studies. Previous studies show that different variant callers generate discordant results for the same genomes (Hwang, et al., 2015; Warden, et al., 2014). But it remains unclear whether using different variant callers ultimately leads to meaningful differences in the set of relevant variants of the interpretation pipelines.

The consequence of a variant can also be transcript specific: a single variant can impact different transcripts of the same gene differently. Therefore, proper transcript choice for a gene when assessing variant impact is important (McCarthy, et al., 2014). Ensembl and APPRIS (Rodriguez, et al., 2013) each provide a dataset of primary transcripts. Ensembl canonical transcripts are defined based on CDS or transcript length, while APPRIS principal isoforms are defined based on the annotations of structure, function and conservation for each transcript. Alternatively, one could consider the impact of the variant in all transcripts of a gene and select one with the most severe impact. This approach, while more sensitive, will be prone to high false positive findings from some minor transcripts. Finally, transcript choice could also be motivated by the biology of the disease, for example by selecting transcripts that are generally expressed in highest levels in tissues that are known to be relevant to the disease.

Population minor allele frequency (MAF) of a variant is another commonly used parameter, typically incorporated in rare genetic disease analysis pipelines to exclude variants whose frequencies are higher in populations than estimated for disease-causing variants according the prevalence of the rare disease. Several large population genetic databases exist, but the MAF for the same variant may differ among these databases, because of the differences in sequencing technology, represented ethnic groups, sequencing quality, and coverage in certain gene regions (Genomes Project, et al., 2015; Lek, et al., 2016). Additionally, MAF thresholds may need to be adjusted according to the prevalence and penetrance of the diseases under consideration (Lek, et al., 2016; Taylor, et al., 2015).

Many computational methods have been developed to predict the deleterious effect of a variant (Hu, et al., 2019). Most tools, like MutPred2 (Pejaver, et al., 2017), MetaSVM (Dong, et al., 2015), REVEL (Ioannidis, et al., 2016) and VEST (Carter, et al., 2013) are focused on missense variants while some other tools also identify impactful splicing variants (Jian, et al., 2014) and noncoding variants (CADD (Kircher, et al., 2014)). To understand whether and how to incorporate predictions from any such tools in an interpretation pipeline requires careful consideration before finalizing a pipeline.

Depending on the goal of the analysis, the optimal values and cutoffs for each of the parameters in analysis pipelines discussed above will differ. For example, for exome analyses in diagnostic settings, the goal of the analysis will be to locate the genetic variants that are consistent with the individual’s clinical features. Such analyses typically emphasize sensitivity over specificity, with appreciation that reviewers will evaluate the variants for significance. Alternatively, if the goal of the analysis is to screen a large number of asymptomatic individuals who may or may not have any genetic disease, the sensitivity and specificity balance may differ. Since the majority of the population is unaffected by rare diseases, even a small reduction in the specificity could lead to a large number of false positives in the population screened. Therefore, the optimal values and thresholds in a pipeline for a diagnostic test may not be applicable for a screening test.

A well-designed pipeline should be robust to small parameter perturbations and yield stable outputs. The design should focus the efforts on refining parameters that matter, not those which yield no changes in perturbation. To study the robustness of analysis pipelines, we developed a framework that started with building initial interpretation pipelines based on prior clinical knowledge and relevant features and then constructed a group of derived pipelines by iteratively perturbing values one parameter at a time from the initial pipelines. This framework enabled us to systematically explore how both the sensitivity and specificity are affected by perturbations to different parameters in the analysis pipelines.

It is challenging to recruit large cohorts of rare disease patients for sequencing. In this study, we applied our perturbation framework to a set of 138 de-identified samples from infants affected with Mendelian inborn errors of metabolism (IEM). Informed by the perturbation analyses in our dataset, we were able to design a robust pipeline that may be appropriate for screening of Mendelian IEMs. This framework of perturbation analysis may be generally applicable and can be informative in other sequencing interpretation pipelines.

## 2 Methods

### 2.1 Datasets, genes and disorders

We sequenced 138 exomes with DNA extracted from the dried blood spots of subjects affected with one of IEMs, initially identified by tandem mass spectrometry by the California Newborn Screening Program (NBSeq affected cases). We also sequenced exomes of 40 cases initially screened positive by MS/MS but determined to be unaffected upon further clinical follow-up (NBSeq unaffected cases). The exome analysis was restricted to a set of IEM genes (Table S1) known to be associated with those disorders.(Adhikari, et al.)

### 2.2 Exome sequencing and analysis

DNA extraction, library preparation and sequencing were carried out following a previously described protocol.(Bassaganyas, et al., 2018) The reads from the NBSeq exomes were mapped to the human reference genome GRCh37 assembly (Aug 2009 release), using BWA-mem v0.7.10 (Li and Durbin, 2009). The resulting BAM files were sorted, indexed and marked for PCR duplicate reads by Picard v0.7.10 (http://picard.sourceforge.net).

### 2.3 Variant calling and annotation

Three different variant callers were run to detect variants in the exomes: GATK 3.3 UnifiedGenotyper (UG), GATK 3.3 HaplotypeCaller (HC) and Platypus 0.8.1. The GATK variant quality scores were recalibrated by VQSR (Variant Quality Score Recalibration), and all 178 samples were jointly called. The annotation of resulting variants was performed using our in-house tool Varant (http://compbio.berkeley.edu/proj/varant/). Variant location region, mutation type, transcript and splice annotations were annotated based on the Gencode annotation (v19). For each of the IEM genes analyzed in the exomes, key transcripts were annotated in four different ways: transcript annotated as canonical transcript according to the Ensembl annotation (https://www.ensembl.org), transcript labeled as a principal isoform according to the APPRIS database (Rodriguez, et al., 2013), transcript where the variant effect is most severe among all alternative transcripts and finally, transcripts with highest expression in liver based on TPM (Transcripts Per Kilobase Million) in GTEx v6 tissue-specific RNA-seq data (Consortium, 2013). Variant minor allele frequency (MAF) was obtained from 1000 genomes (phase 3) (Genomes Project, et al., 2015), NHLBI Exome Sequencing Project (ESP, ESP6500SI-V2-SSA137 dataset) (Tabor, et al., 2014), and Exome Aggregation Consortium (ExAC v 0.3.1)(Lek, et al., 2016). Variants were annotated with effect predictions from CADD v1.3 (Kircher, et al., 2014), MetaSVM from dbNSFP v3.1a(Dong, et al., 2015), RF score from dbscSNV v1.0 (Jian, et al., 2014) and loss of function (LoF) annotation from LOFTEE tool [v0.2] (https://github.com/konradjk/loftee). Disease associated variants were annotated from HGMD v2014.1 (Stenson, et al., 2014) and Clinvar (dataset ftp://ftp.ncbi.nlm.nih.gov/pub/clinvar/tab_delimited/variant_summary.txt.gz accessed 24th April 2016). Specific filtering thresholds for each annotation feature are shown in table S2.

### 2.4 Variant interpretation pipelines

The pipeline development framework perturbs and updates an initial reference pipeline until reaching a stable pipeline. Newborn screening of IEMs requires both high sensitivity and high specificity. To design an effective exome analysis pipeline that meets the requirements for screening, we started from two initial reference pipelines, pipeline A-1 (favoring sensitivity) and pipeline B-1 (favoring specificity) (Table 1). The two pipelines had the same architecture (Figure 2A). Potentially disease-causing variants were flagged by the pipelines through two primary filtering arms. The first arm reported variants that are possibly pathogenic based on their predicted impact and rarity, while the second arm reported variants that are curated as pathogenic in disease databases (HGMD and Clinvar). Among all reported variants for an individual exome in the gene list, the pipeline then reported the genes where ≥1 homozygous or hemizygous variant or ≥ 2 heterozygous variants were reported. To identify which commonly used parameters in exome analysis pipelines would influence the sensitivity and specificity, we systematically explored the following parameters: IEM gene list, variant callers, choices of transcript models, choice of population databases, minor allele frequency (MAF) thresholds in population databases, databases of predicted and curated variants, and choice of inheritance models. We studied the impact of each parameter on overall performance by altering a single parameter or a few parameters at a time (Table S2). Based on the results from the perturbation analysis, we finally developed a tuned pipeline (pipeline C-1) with balanced sensitivity and specificity (Table 1).

**Figure 1.**
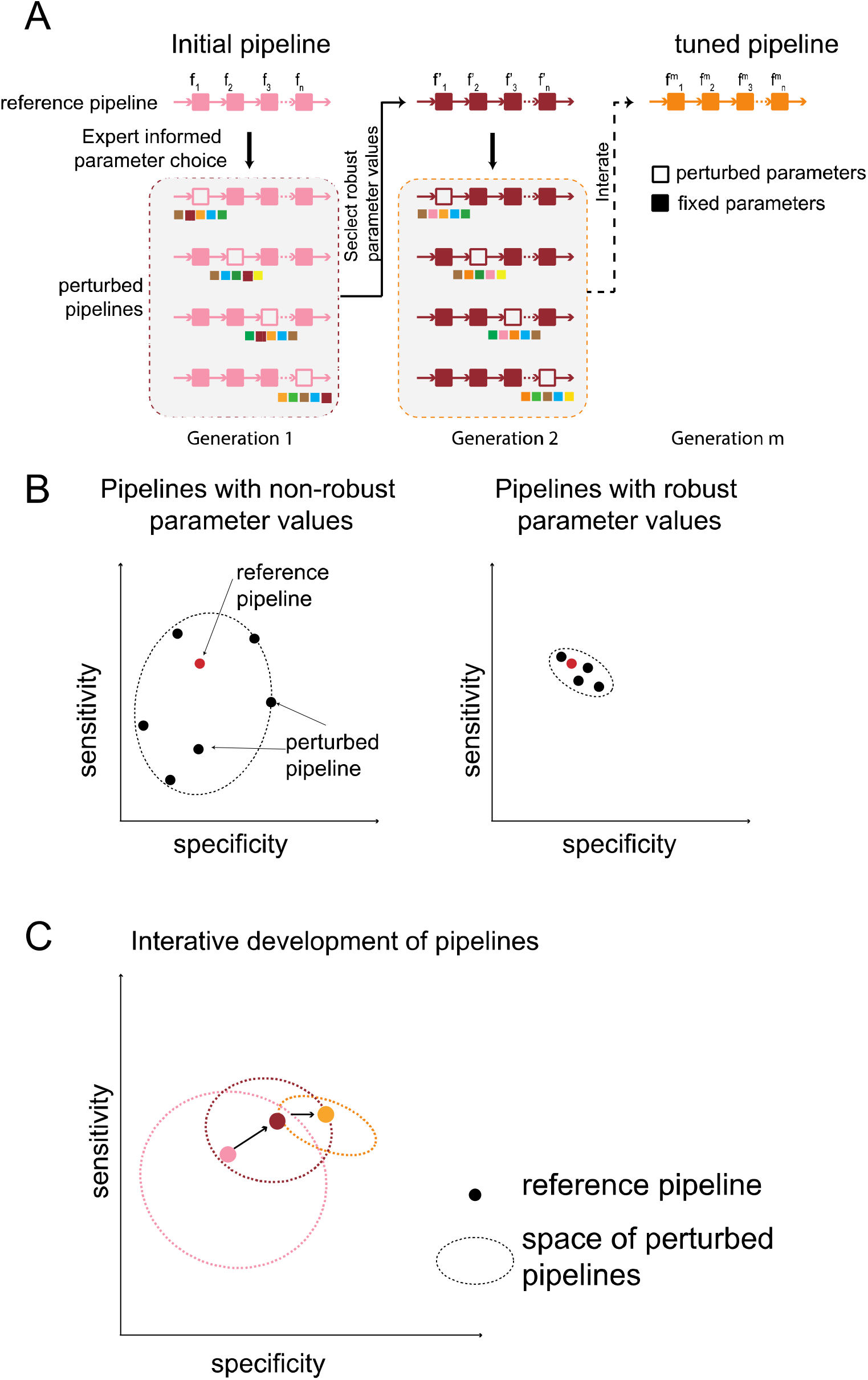
Framework of perturbation analysis. **A)** An initial pipeline has multiple parameters which then are perturbed one by one to generate a group of derived pipelines. Robust parameters are identified by comparing the results from the derived pipelines. This information guides an updated reference pipeline design and based on the reference to perform a new round perturbation analysis. **B)** Sensitive parameters have large impact on pipeline performance, present varied sensitivity and specificity. Pipeline using robust paraments should be clustered close to the reference pipeline in terms of sensitivity and specificity. **C)** Starting from two initial pipelines, pipelines were updated using the perturbation framework till reaching a final pipeline with optimal sensitivity and specificity.

**Figure 2.**
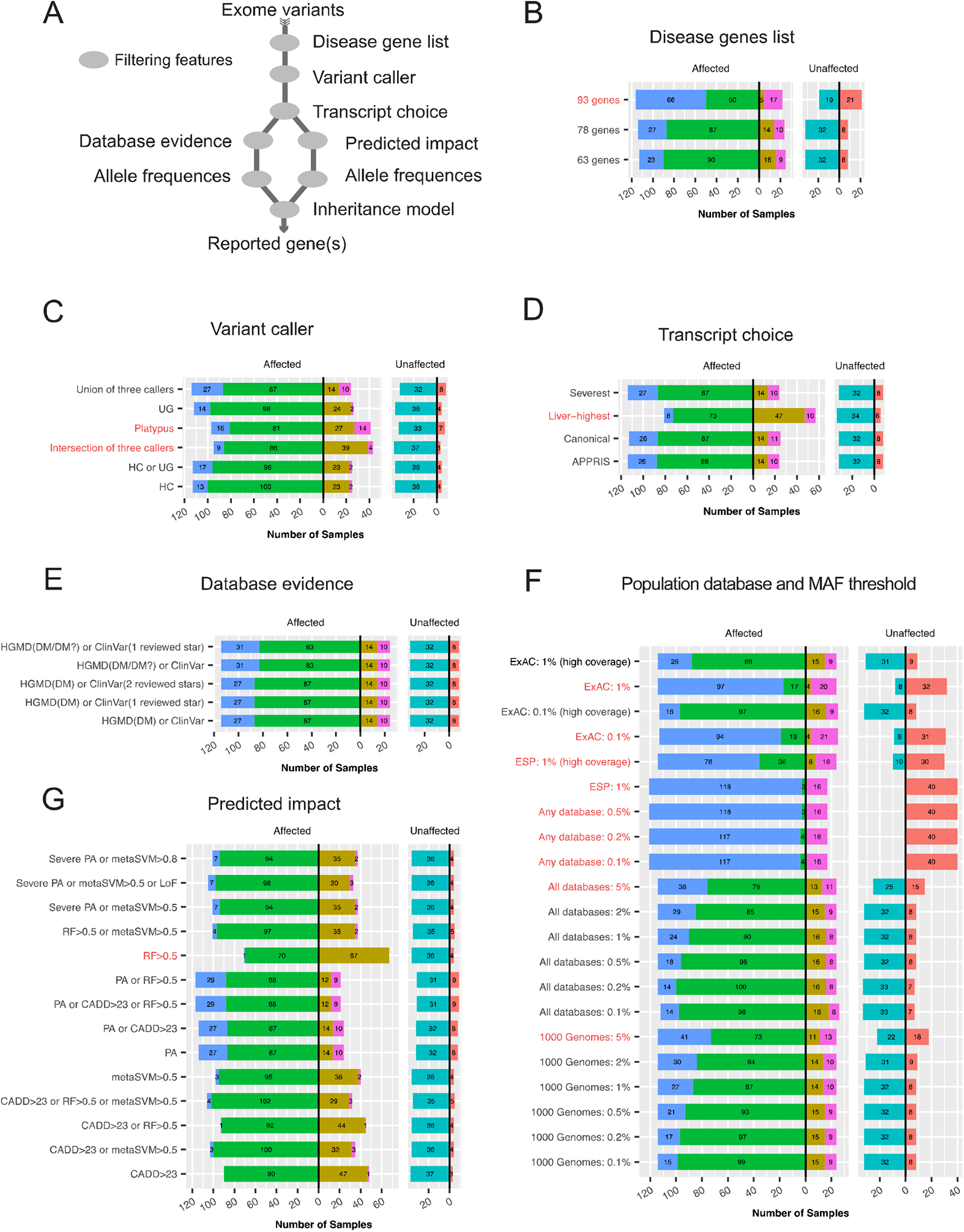
Pipeline design and perturbation of group A pipeline. **A)** The variant interpretation pipeline has a two-arm architecture. Variants from each individuals’ exome were filtered through multiple layer of features and finally output disease genes where at least one homozygous variant or two heterozygous variants passed the filters. Non-robust parameters (red text) of these filter features **B), C), D), E)** and **F)** were identified through assessing the performance of group A pipelines.

**Table 1.**
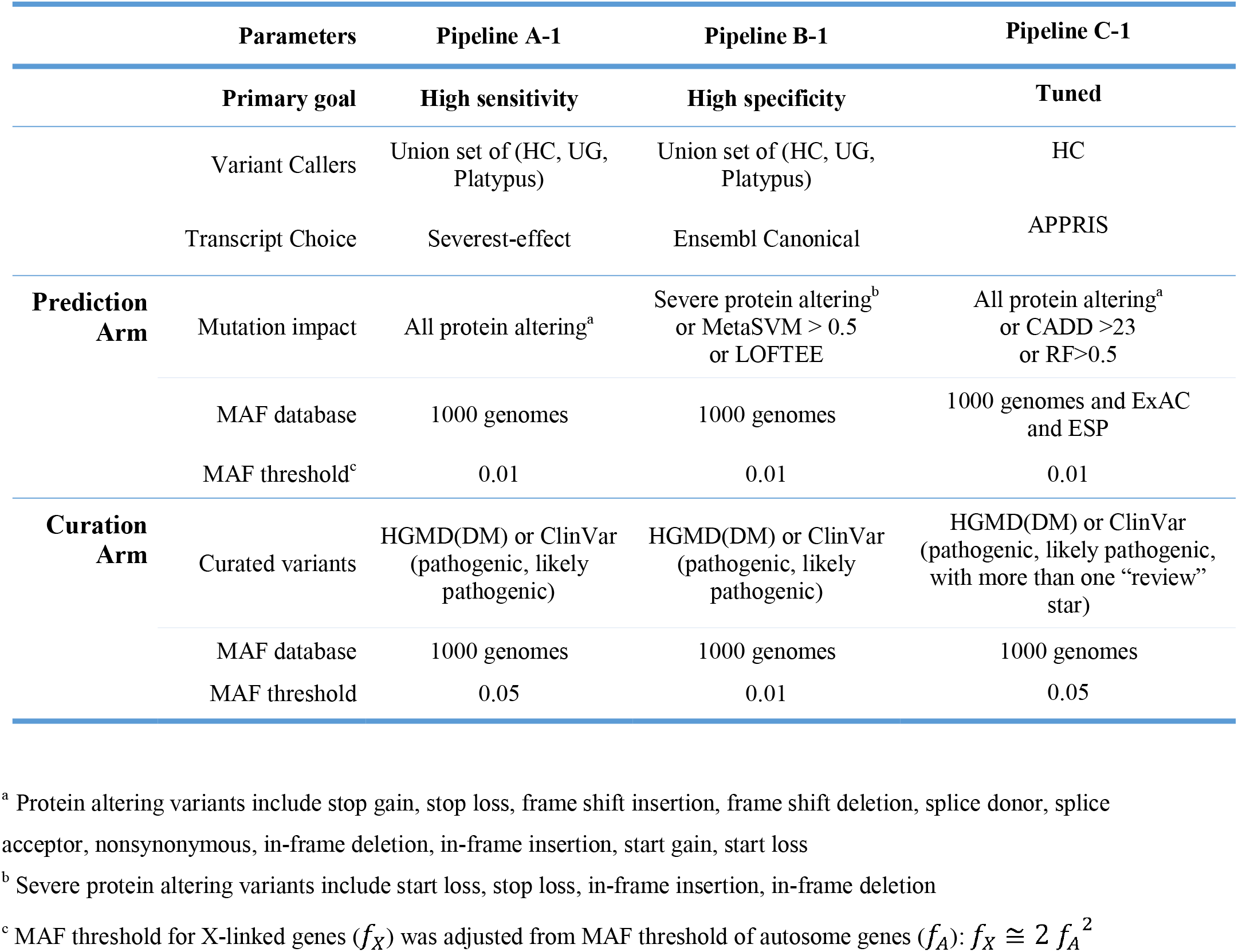
Parameters of reference pipelines.

|  | Parameters | Pipeline A-1 | Pipeline B-1 | Pipeline C-1 |
| --- | --- | --- | --- | --- |
|  | Primary goal | High sensitivity | High specificity | Tuned |
|  | Variant Callers | Union set of (HC, UG, Platypus) | Union set of (HC, UG, Platypus) | HC |
|  | Transcript Choice | Severest-effect | Ensembl Canonical | APPRIS |
| **Prediction Arm** | Mutation impact | All protein altering<sup>a</sup> | Severe protein altering<sup>b</sup> or MetaSVM > 0.5 or LOFTEE | All protein altering<sup>a</sup> or CADD >23 or RF>0.5 |
|  | MAF database | 1000 genomes | 1000 genomes | 1000 genomes and ExAC and ESP |
|  | MAF threshold<sup>c</sup> | 0.01 | 0.01 | 0.01 |
| **Curation Arm** | Curated variants | HGMD(DM) or ClinVar (pathogenic, likely pathogenic) | HGMD(DM) or ClinVar (pathogenic, likely pathogenic) | HGMD(DM) or ClinVar (pathogenic, likely pathogenic, with more than one “review” star) |
|  | MAF database | 1000 genomes | 1000 genomes | 1000 genomes |
|  | MAF threshold | 0.05 | 0.01 | 0.05 |

<sup>a</sup> Protein altering variants include stop gain, stop loss, frame shift insertion, frame shift deletion, splice donor, splice acceptor, nonsynonymous, in-frame deletion, in-frame insertion, start gain, start loss
<sup>b</sup> Severe protein altering variants include start loss, stop loss, in-frame insertion, in-frame deletion
<sup>c</sup> MAF threshold for X-linked genes ( $f_X$ ) was adjusted from MAF threshold of autosome genes ( $f_A$ ): $f_X \cong 2 f_A^2$

### 2.5 Assessment of pipeline predictions

The set of output genes calculated by a pipeline for each NBSeq individual’s exome was compared to the individual’s disorder based on a gene-to-disorder mapping dictionary. The affected cases where all the output gene(s) from the pipeline were consistent with the individual’s disorder were labeled true positive (TP). Affected cases with no output genes(s) from the pipeline were labeled false negative (FN). If all output gene(s) were inconsistent with the individual’s disorder, the case was labeled (FN&FP), and if only a subset of the output genes were consistent with the individual’s disorder, the case was labeled as (TP&FP). We used the true positive rate (the ratio of TP cases among total affected cases) and positive rate (the ratio of TP and TP&FP cases among affected cases) to calculate the sensitivity of each pipeline. For unaffected individuals, the pipeline outputs were grouped into two categories: 1) True negative (TN): cases for which none of the IEM genes was flagged in the individuals; 2) False positive (FP): cases for which the pipeline flagged one or more of the IEM genes in the individuals. The specificity of a pipeline was defined as the fraction of TN predictions among unaffected individuals (N=40). We also assessed the pipeline’s specificity on 2504 individuals from the 1000 genomes project. The possibility of these individuals’ having any of these metabolic disorders should be lower than 1 per 1000, based on the prevalence of the disorders. Thus, the specificity was calculated as the ratio of number of individuals flagged for any of the IEM genes by the pipeline, against the 2504 individuals.

To ensure an unbiased design of the pipelines and an objective interpretation of the results, the researchers who developed the analysis pipelines and assessed the results (YW, ANA, SEB) were blinded to the diagnosis of individual cases. A Perl script was developed to perform the automated assessment of the pipelines (https://github.com/nbseq1200/NBSeq1200paper/tree/master/assessment). The assessment tool was run by an independent researcher (MNK) against the case-by-case diagnosis data provided by RJC and RCG.

## 3. Results

### 3.1. Pipeline perturbation framework

We developed a framework to iteratively perturb and update parameters to build robust, optimized variant interpretation pipelines (Figure 1). From an initial reference pipeline using parameter filters typical in diagnostic exome analyses of rare Mendelian disorders, the framework perturbs one or a few parameters to obtain a group of “perturbed pipelines”. The perturbed parameter values are chosen based on prior knowledge about the disorders under study. By comparing the outputs from the derived pipelines to that of the initial reference pipeline, parameters that generally are robust to perturbations can be identified (Figure 1A). Robust parameters when perturbed should generally have small impact on the pipeline’s sensitivity and specificity (Figure 1B). A new reference pipeline is then obtained by updating the initial reference pipeline with robust parameters values that locally optimize both sensitivity and specificity (Figure 1C). This process of updating the parameters is repeated in multiple rounds until both the sensitivity and specificity of the pipelines remain stable to perturbations.

To demonstrate the utility of this framework, we applied it to guide pipeline design for the UCSF NBSeq project, for which the goal was to evaluate the utility of exome sequencing as a potential primary screen for rare IEMs. We developed screening variant interpretation pipelines considering commonly used parameters, including disease gene list, variant callers, transcripts, predicted variant impact, disease database annotations, and minor allele frequencies (Figure 2A). Unlike diagnostic exomes where sensitivity is generally prioritized, a newborn screen applied on a population-scale requires both very high sensitivity and specificity. Therefore, we designed two initial reference pipelines with initial parameter choices as described in Table 1 that prioritized either sensitivity (A-1) or specificity (B-1). Two groups of derived pipelines were then obtained following the approach described above. After fixing the robust parameters, the performance of the pipelines in the two groups were converged to a similar set of parameter choices, which then guided the design of a final pipeline (C-1).

### 3.2 Choice of gene list

Expanding the gene list to more candidate genes could potentially identify more affected cases, but would also increase the likelihood of the false positive findings through spurious pathogenicity calls. As shown in (Figure 2B), the pipeline identified one more affected case correctly without incurring false positives when the exome analysis was expanded from the conservative gene list of 63 most confident genes associated with IEMs under study to additional 15 IEM-related genes (78 gene list). However, upon expanding the gene list even further to include more candidate IEM-related genes (93 gene list), only 2 additional affected cases were correctly identified but at the cost of simultaneously increasing 13 false positives. Therefore, the 78 gene list was chosen for the final pipeline.

### 3.3. Choice of variant callers impacts prediction results

To assess the impact of using different variant callers, we considered variant calls generated by individual tools or combined from multiple tools. Three variant callers were run: UnifiedGenotyper (UG), HaplotypeCaller (HC) from GATK, and Platypus. Using either UG or HC alone missed 26 and 25 cases respectively, and both had 4 false positive cases (Figure 2C) UG and HC gave similar results in group B pipelines (Figure S1B). However, using Platypus individually missed more affected cases in group A (41 cases) and had more false positive cases (7 cases). The union set of variants called by UG and HC gave similar results compared to the results by using either tool individually (Figure 2C and Figure S1B). But combining the results from Platypus worsened the pipelines’ performance on our dataset: using a union set of variants from the three tools had a higher false positive rate; while using an intersection set of variants from the three callers missed many more cases (Figure 2C). We further manually checked some of discrepant cases among Platypus, HC and UG. The sequencing reads supported the results from HC and UG. Based on these results, we chose HC for the final pipeline.

### 3.4. Prediction results are generally insensitive to transcript choice

Almost all multi-exon human genes can produce multiple RNA transcripts (Pan, et al., 2008; Wang, et al., 2008). We expected that considering a variant’s severest impact on all possible transcript (severest-impact transcript) would be more sensitive than only considering the impact on the primary transcript. To test this hypothesis, we compared performance of pipelines that considered two different standard human transcript sets of the IEM genes (Figure 2D). Ensembl (Aken, et al., 2017) and APPRIS (Rodriguez, et al., 2013) each offer a set of primary transcripts. The difference in overall performance for all the transcript sets considered was small (considering the ‘severest effect’ transcript identified one more affected case than considering canonical transcript, and one more TP&FP case than using APPRIS principal transcript), but revealed two interesting cases. In one case, an affected individual could be identified using the “severest effect” transcript, but would have been missed if only the Ensembl canonical transcript was used. This individual had a start gain variant in gene *SLC22A5* based on a non-canonical transcript which was the APPRIS principal transcript (ENST00000245407). However, considering the severest transcript also contributed to a false positive call of gene *ETFB* in one case, where a homozygous variant was flagged because it was an in-frame insertion in the Ensembl canonical transcript (ENST00000354232). However, this variant was located in the intron of the APPRIS principal transcript (ENST00000309244). In both *SLC22A5* and *ETFB*, the APPRIS transcripts are conserved in multiple species while the canonical transcripts are human specific (according to the APPRIS database). Thus, considering APPRIS transcripts was reasonable in these two cases and also gave better overall performance on the NBSeq data set. As metabolic genes mainly operate in liver, we also considered using the transcript with the highest expression in liver. We selected the most abundant transcript in liver of each IEM gene based on the liver expression data from GTEx (v6). However, using this transcript set led to worse prediction results (Figure. 2D). For some genes, the most abundant transcript in liver had an incomplete CDS or was predicted to be a target of nonsense-mediated RNA decay (Table S3). Thus, it is not always appropriate to choose the transcript for assessing variant impacts based solely on abundance of expression.

### 3.5. Inclusion of likely-pathogenic curated variants had minor impact

Choices of curated variant pathogenicity evidence from HGMD or ClinVar databases had a minor impact on overall performance. HGMD has two levels of confidence for labeling disease-causing mutations (DM for disease-causing mutations and DM? for likely disease-causing mutations). ClinVar uses number of “review” stars to indicate confidence of the variant pathogenicity (one star for single assertion and two stars for multiple assertions with no conflicting assertion). However, pipelines which considered more stringent ClinVar evidence (requiring at least one star or two stars) did not change the results (Figure 2E and Figure S1D). While pipelines that incorporated DM? variants did not reduce the number of missed cases, they increased the number of true and false positives by flagging additional incorrect gene(s) besides the correct disease gene (Figure 2E).

### 3.6. Pipeline results are sensitive to variant impact prediction methods

We explored choices of different combinations of computational variant effect prediction tools and evaluated their performance on correctly implicating the underlying disorders in the affected individuals. Our initial pipelines had two filter arms (Figure 2A): the curation arm which considered variants from curated disease databases, and the prediction arm which considered variants flagged by prediction tools (see details in methods). To untangle the possible interactions of parameters from the curation and prediction arms, we first tested a set of variant impact prediction tools utilizing a group of one-arm pipelines retaining only the prediction arms. Removal of the curation arm led to varying results based on the prediction tools used (Figure S2). For example, pipelines which used CADD and metaSVM alone, identified 52 and 71 TP cases respectively. The pipelines considering union sets of deleterious variants flagged by both CADD and metaSVM, identified 83 TP cases. In pipelines with both filtering arms, the difference in results from using different tools was smaller (Figure 2G). With the curation arm in place, there were 90, 95 and 100 TP cases identified by CADD, metaSVM and combination of the two tools, respectively.

Considering all the protein altering (PA) variants (Table 1) identified relatively more positive cases (TP and TP&FP) than using only other tested tools for both group A and group B pipelines (Figure 2G and Figure S1E). Our list of protein altering variants included splice donors and splice acceptors, but didn’t include other exonic or intronic splice impacting variants. The perturbation results showed that incorporating splicing impact prediction (using RF score) improved results for both sets of pipelines (Figure. 2G and Figure S1E) by increasing the number of TP cases.

### 3.7. MAF database and threshold choice yield dramatic differences on prediction results

Because IEMs are rare in newborns, the variant frequency in reference populations is an important feature for identifying disease-causing variants. Choice of different reference population databases and MAF thresholds yielded dramatic differences in pipeline results (Figure 2F and S2F). For example, in group A pipelines, we used MAFs obtained from the 1000 genomes project, and explored frequencies in the range 0.1% to 5% gradually. The results showed that higher MAF thresholds allowed the pipeline to flag more variants, leading to higher numbers of TP & FP cases (from 15 to 41 cases) but did not help reduce the number of missed cases (all pipelines missed 24 affected cases). The pattern of increasing TP & FP cases, without an accompanying decrease in the number of missed cases was independent of the source of MAF values.

Prediction results were also surprisingly sensitive to database choice. Using MAF from the NHLBI Exome Sequencing Project (ESP) alone lead to more incorrect gene calls in affected individuals and more false positive results in unaffected individuals, compared to using MAF from 1000 genomes. Choosing MAF from the Exome Aggregation Consortium (ExAC) database was slightly worse, compared to using the same MAF threshold from 1000 genomes. To avoid any sequencing coverage bias, the pipeline only considered variants in well covered regions in ESP and ExAC exomes. Ignoring coverage information, caused the performance of these pipelines to be much worse. However, we also found common variants with MAF larger than 1% in 1000 genomes to be reported by ESP as very low MAF even when the coverage was good.

### 3.8 Perturbation analysis informs the design of the tuned pipeline

Based on results from previous perturbation analyses, we identified robust values for parameters that were suitable for an exome screening pipeline and designed a tuned pipeline C-1 (Table 1). The tuned pipeline was more sensitive and specific compared to pipelines A-1 and B-1 (Figure 3). The sensitivity was calculated as the fraction of TP cases, in which only the correct IEM gene(s) were flagged, among all affected cases (Figure 3). The specificity was calculated based on the fraction of TN cases among unaffected cases.

**Figure 3.**
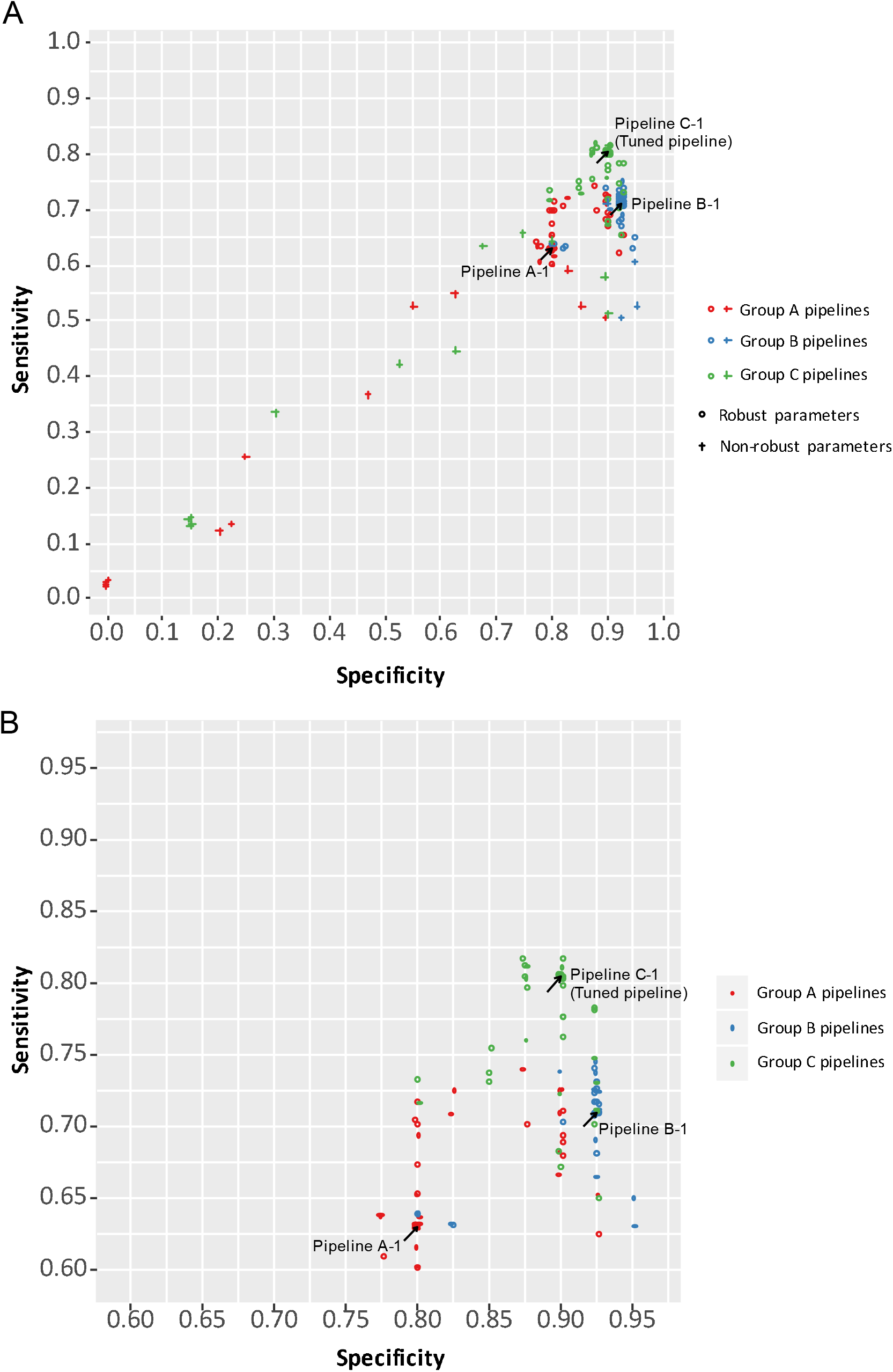
Performance of group A, B and C pipelines. Sensitivity of each pipeline was calculated as the fraction of true positive cases among affected individuals. Specificity was calculated as the fraction of true negative cases among unaffected individuals. **A)** The performance of all perturbed pipelines from each group. **B)** After excluding pipelines with non-robust parameters, the convex hull was drawn to represent the sensitivity-specificity space of each group of pipelines.

To test the robustness of the tuned pipeline C-1, we performed similar perturbation analyses by changing the value of one parameter at a time of the tuned pipeline to generate group C pipelines. Most group C pipelines clustered to C-1, except pipelines with non-robust parameters which had been identified by perturbation analysis of Group A and B pipelines (Figure 3A). After excluding the non-robust parameters (red text in Figure 2), the group C pipelines have higher sensitivity and specificity compared to group A and B pipelines (Figure 3B).

## 4. Discussion

### 4.1. Findings from the perturbation analysis

The perturbation framework that changes one parameter at a time from an initial reference pipeline and compares the performance of all generated pipelines efficiently filtered out non-robust parameters. The perturbation analyses showed that choice of variant callers, variant impact prediction tools, MAF threshold and MAF databases have large impacts on pipeline performance. Transcript isoform choice and curated variant evidence were found to be relatively unimportant for identifying disease-causing variants in IEM genes. The perturbation analyses also revealed that a two-arm structure that integrated curated and predicted parameters was more robust than one that included either arm alone.

### 4.2. Limitations of this study

Our perturbation analysis considered commonly used exome parameters to build a systematic framework to study the impact of those parameters. However, this study has limitations in the following aspects. First, it did not explore the whole combinatorial parameter space. We assessed the pipeline performance on a limited set of 178 exomes and some of the disorders were too rare to provide enough training samples. Because of the limited sample size, we used a blinded approach instead of cross validation to estimate the sensitivity and specificity. In Figure 3, we calculated the sensitivity by only considering the TP cases. Another way to estimate the sensitivity would be to calculate the fraction of all positive cases in which at least one correct IEM was flagged (TP and TP&FP cases). Group C pipelines also showed better overall performance than Group A and B pipelines (Figure S3). We calculated the specificity from the 40 unaffected cases. These cases may not be a good representation of the real-world unaffected population: notably, they were false positive results from the initial MS/MS screening. However, the specificity estimated from this dataset correlated well with the specificity estimated from running the same pipeline on the 1000 Genomes data (Figure S3). Third, the assessment criteria were based on the requirements of screening, and the specific parameters chosen by the tuned pipeline were only suited for screening this group of rare monogenetic metabolic disorders. To find the optimal parameters for other applications, the full set of perturbation analyses would have to be run.

### 4.3. General application of this framework

The work described in this paper addresses the challenge of developing a robust pipeline for clinical interpretation of genetic variation that simultaneously achieves high sensitivity and specificity, especially when the data available for model building is limited. We use our previous experience in identifying causal variants of rare inherited disorders and our experience in organizing and participating in a community-wide assessment of methods for genome interpretation (CAGI) (Andreoletti, et al., 2019) to guide our selection of a set of parameters that are likely to be important in identifying genetic variations that cause inherited disorders. With the parameters in hand, and again guided by our prior experience, we chose appropriate values for these parameters to develop two pipelines – one that prioritizes sensitivity and another that prioritizes specificity. Having established the two base-line pipelines we undertook a systematic exploration of parameter values to develop a pipeline that balances the twin requirements of sensitivity and specificity. Such an analysis identified robust parameters along with appropriate thresholds resulting in a pipeline well suited to the purpose.

Our approach is not without limitations. The developed pipeline cannot be asserted to be optimal. We however note that obtaining a highly optimized pipeline with limited data, as is commonly aimed for in machine learning approaches, may not even be desirable. Further, it is quite possible that the order in which the parameters are tuned can have a bearing on the final pipeline. However, as our results show, for the problem at hand a robust pipeline was achieved. We postulate that a similar approach could apply to other problems, where underlying characteristics of the problem are known (for e.g. disease prevalence and causal genes in our case), the available data for learning a pipeline is limited, and human expertise in the problem domain is available.

#### Ethics approval and consent to participate

This study was approved by the California Committee for the Protection of Human Subjects (project number 14-07-1650). Residual, de-identified newborn dried blood spot samples were used to prepare DNA for exome sequencing. No sample or DNA remained after WES. If other researchers desire access to data or DBS, they would need to make a separate application to the CDPH (https://www.cdph.ca.gov/Programs/CFH/DGDS/Pages/cbp/default.aspx). California law precludes sharing specimens or uploading individual data derived from them into any genomic data repository.

## Acknowledgements

We thank John-Marc Chandonia, Jingqi Chen, Zhiqiang Hu and Andrew Sharo for insightful discussions about this research. The work was funded by the National Institute of Health grant U19HD077627 as part of the NSIGHT Project, a joint program between the National Human Genome Research Institute and the Eunice Kennedy Shriver National Institute of Child Health and Human Development, NIH. This work was also supported by a research agreement with Tata Consultancy Services. The biospecimens and/or data used in this study were obtained from the California Biobank Program, (SIS request number 496). The California Department of Public Health is not responsible for the results or conclusions drawn by the authors of this publication.

## Author contributions

Y.W, A.N.A and S.E.B. conceived and designed the study. A.N.A., R.J.C, M.K., U.S., P-Y.K., J.M.P. acquired data. Y.W., A.N.A., R.J.C, R.C.G., M.K., J.M.P., R.S., U.S., S.E.B. analyzed data. A.N.A., Y.W., S.E.B. interpreted data. A.N.A., Y.W., U.S. created software. Y.W. wrote the first draft of the manuscript. Y.W., A.N.A., M.K., N.J.R., B.A.K., P-Y.K., J.M.P, R.S., S.E.B. provided critical revisions. S.E.B. was unable to review the final manuscript due to injury.

## Competing interests

Aashish Adhikari is currently an employee of Illumina, Inc. Uma Sunderam and Rajgopal Srinivasan are employees of Tata Consultancy Services (TCS). Jennifer Puck is the spouse of Robert Nussbaum, an employee of Invitae. Steven E. Brenner receives support at the University of California Berkeley from a research agreement from TCS.

## Supplementary Tables

Table S1 IEM gene list

Table S2 Parameters of group A, B and tuned pipelines

Table S3 Performance of group A, B and tuned pipelines based on NBSeq data

Table S4 Most abundant transcript in liver

## Supplementary Figures

**Figure S1 Performance of group B pipelines**

**Figure S2 Performance of one-arm pipelines.** A) Group A, curated arm pipeline. B) Group B, curated arm pipeline. C) Group A predicted impact arm pipeline. D) Group B predicted impact arm pipeline.

**Figure S3 Performance of group A, B and C pipelines.** Sensitivity of each pipeline was calculated as the fraction of all positive (TP and TP&FP) cases among affected individuals. Specificity was calculated as the fraction of true negative cases among unaffected individuals. **A)** The performance of all perturbed pipelines from each group. **B)** After excluding pipelines with non-robust parameters, the convex hull was drawn to represent the sensitivity-specificity space of each group of pipelines.

**Figure S4. Comparison of pipeline specificities estimated from NBSeq and 1000 Genomes data.** The linear regression lines for group A and B pipelines are in red and blue respectively. The black dashed line showed the Y=X. Data points below this line suggested NBSeq data gave lower estimation of specificities than those from 1000 Genome data.

## References

Adhikari, A.N., et al. The Role of Exome Sequencing in Newborn Screening for Inborn Errors of Metabolism. Nat Medicine (in press).

Aken, B.L., et al. Ensembl 2017. Nucleic acids research 2017;45(D1):D635–D642.

Bassaganyas, L., et al. Whole exome and whole genome sequencing with dried blood spot DNA without whole genome amplification. Hum Mutat 2018;39(1):167–171.

Carter, H., et al. Identifying Mendelian disease genes with the variant effect scoring tool. BMC Genomics 2013;14 Suppl 3:S3.

Consortium, G.T. The Genotype-Tissue Expression (GTEx) project. Nature genetics 2013;45(6):580–585.

Cummings, B.B., et al. Transcript expression-aware annotation improves rare variant interpretation. Nature 2020;581(7809):452–458.

Dong, C., et al. Comparison and integration of deleteriousness prediction methods for nonsynonymous SNVs in whole exome sequencing studies. Human molecular genetics 2015;24(8):2125–2137.

Genomes Project, C., et al. A global reference for human genetic variation. Nature 2015;526(7571):68–74.

Hu, Z., et al. VIPdb, a genetic Variant Impact Predictor Database. Hum Mutat 2019;40(9):1202–1214.

Hwang, S., et al. Systematic comparison of variant calling pipelines using gold standard personal exome variants. Scientific reports 2015;5:17875.

Ioannidis, N.M., et al. REVEL: An Ensemble Method for Predicting the Pathogenicity of Rare Missense Variants. Am J Hum Genet 2016;99(4):877–885.

Jian, X., Boerwinkle, E. and Liu, X. In silico prediction of splice-altering single nucleotide variants in the human genome. Nucleic acids research 2014;42(22):13534–13544.

Kircher, M., et al. A general framework for estimating the relative pathogenicity of human genetic variants. Nature genetics 2014;46(3):310–315.

Lek, M., et al. Analysis of protein-coding genetic variation in 60,706 humans. Nature 2016;536(7616):285–291.

Li, H. and Durbin, R. Fast and accurate short read alignment with Burrows-Wheeler transform. Bioinformatics 2009;25(14):1754–1760.

McCarthy, D.J., et al. Choice of transcripts and software has a large effect on variant annotation. Genome medicine 2014;6(3):26.

Pan, Q., et al. Deep surveying of alternative splicing complexity in the human transcriptome by high-throughput sequencing. Nature genetics 2008;40(12):1413–1415.

Pejaver, V., et al. MutPred2: inferring the molecular and phenotypic impact of amino acid variants. bioRxiv 2017:134981.

Rodriguez, J.M., et al. APPRIS: annotation of principal and alternative splice isoforms. Nucleic acids research 2013;41(Database issue):D110–117.

Schnappauf, O. and Aksentijevich, I. Current and future advances in genetic testing in systemic autoinflammatory diseases. Rheumatology (Oxford) 2019;58(Supplement_6):vi44–vi55.

Stenson, P.D., et al. The Human Gene Mutation Database: building a comprehensive mutation repository for clinical and molecular genetics, diagnostic testing and personalized genomic medicine. Hum Genet 2014;133:1–9.

Tabor, H.K., et al. Pathogenic variants for Mendelian and complex traits in exomes of 6,517 European and African Americans: implications for the return of incidental results. Am J Hum Genet 2014;95(2):183–193.

Taylor, J.C., et al. Factors influencing success of clinical genome sequencing across a broad spectrum of disorders. Nature genetics 2015;47(7):717–726.

Wang, E.T., et al. Alternative isoform regulation in human tissue transcriptomes. Nature 2008;456(7221):470–476.

Warden, C.D., et al. Detailed comparison of two popular variant calling packages for exome and targeted exon studies. PeerJ 2014;2:e600.

Wright, C.F., FitzPatrick, D.R. and Firth, H.V. Paediatric genomics: diagnosing rare disease in children. Nat Rev Genet 2018;19(5):325.

