## Supplemental Figures for "Perturbation robustness analyses reveal important parameters in variant interpretation pipelines"

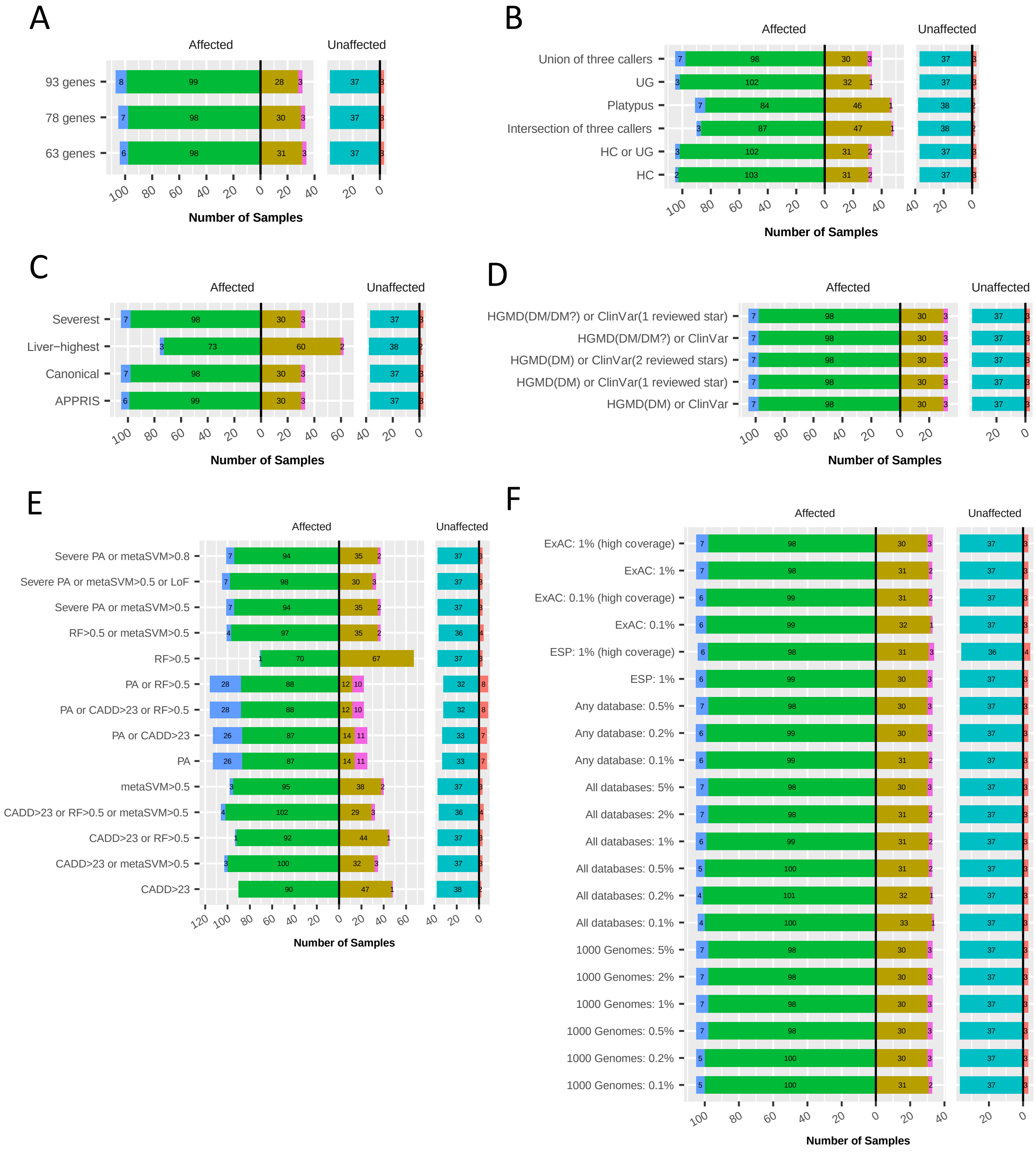


### Figure S1 Performance of group B pipelines

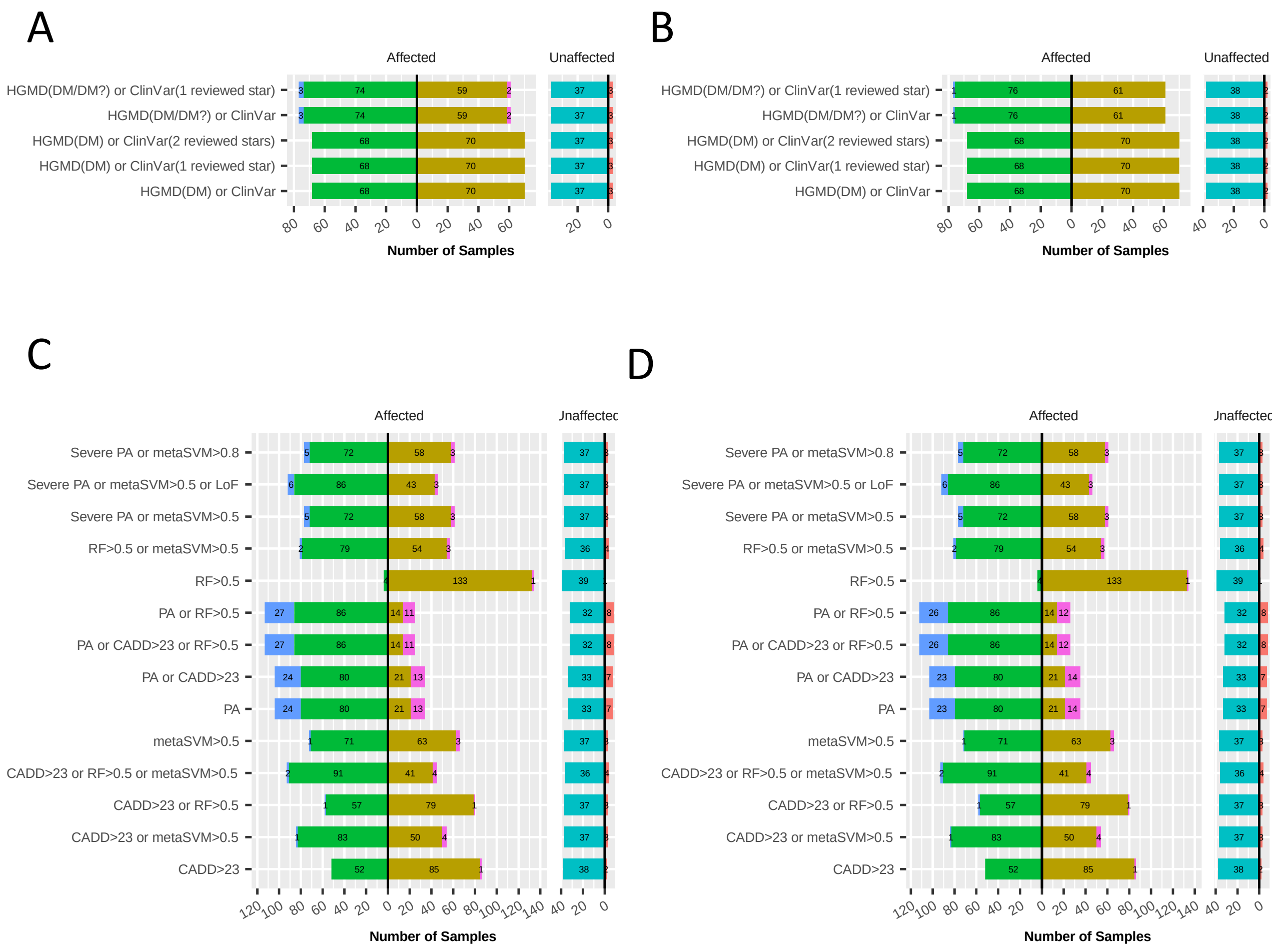


**Figure S2 Performance of one-arm pipelines.** A) Group A, curated arm pipeline. B)Group B, curated arm pipeline. C) Group A, predicted impact arm pipeline. D) Group B predicted impact arm pipeline.

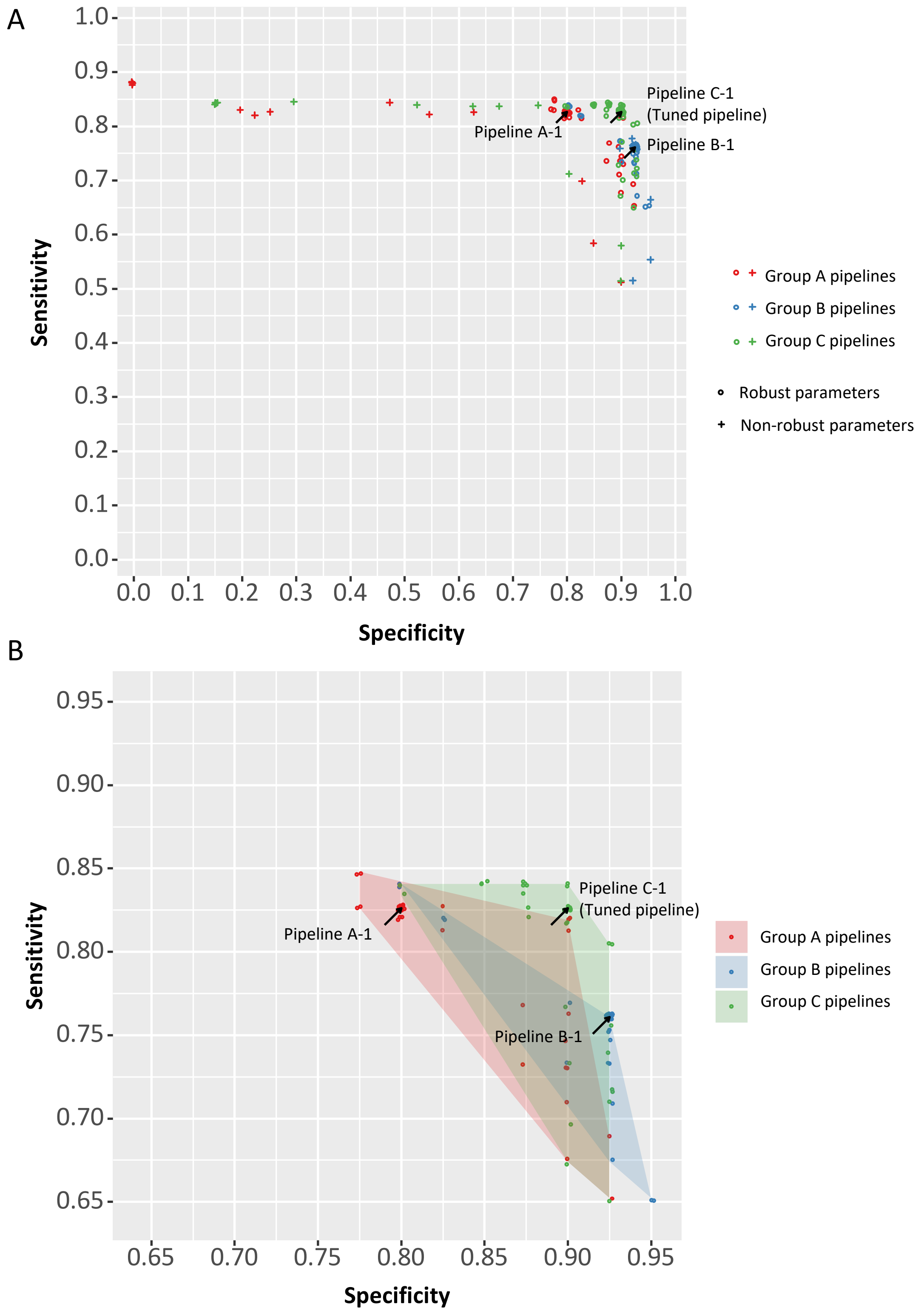


**Figure S3 Performance of group A, B and C pipelines.** Sensitivity of each pipeline was calculated as the fraction of all positive (TP and TP&FP) cases among affected individuals. Specificity was calculated as the fraction of true negative cases among unaffected individuals. **A)** The performance of all perturbed pipelines from each group. **B)** After excluding pipelines with non-robust parameters, the convex hull was drawn to represent the sensitivity-specificity space of each group of pipelines.

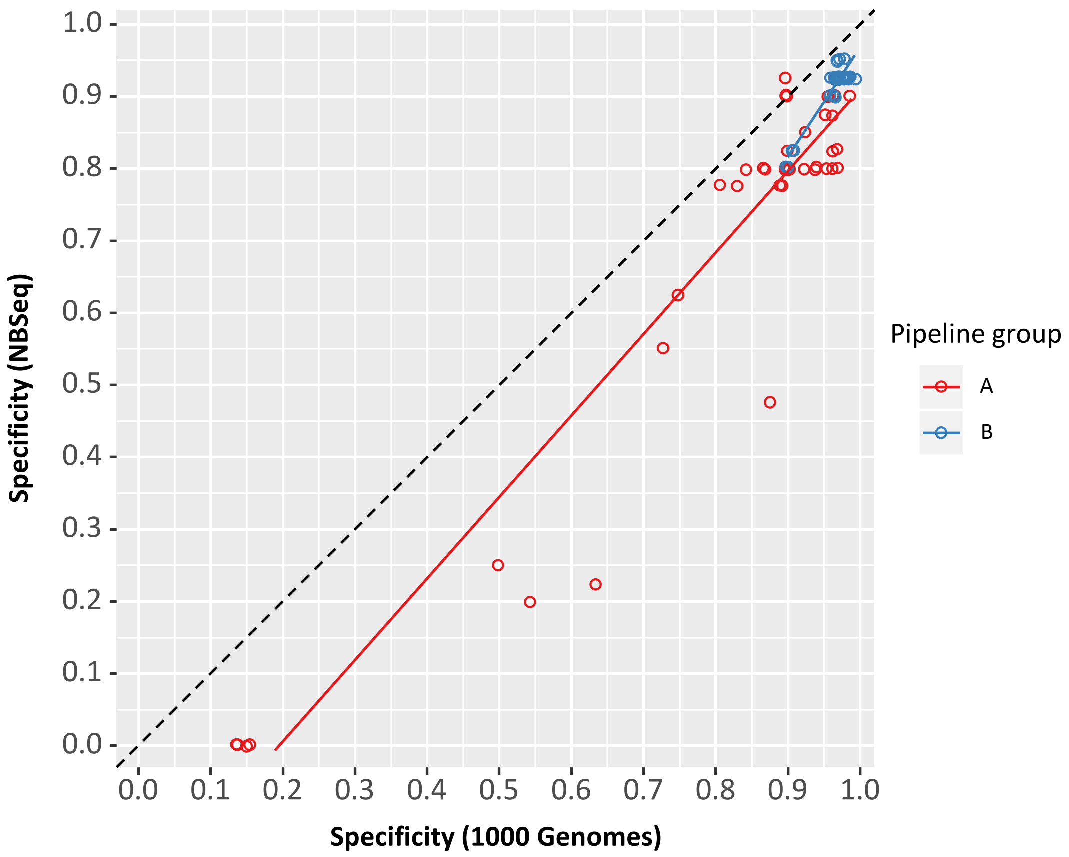


**Figure S4. Comparison of pipeline specificities estimated from NBSeq and 1000 Genomes data.** The linear regression lines for group A and B pipelines are in red and blue respectively. The black dashed line showed the Y=X. Data points below this line suggested NBSeq data gave lower estimation of specificities than those from 1000 Genome data.
